# Population size history from short genomic scaffolds: how short is too short?

**DOI:** 10.1101/382036

**Authors:** Graham Gower, Jono Tuke, AB Rohrlach, Julien Soubrier, Bastien Llamas, Nigel Bean, Alan Cooper

## Abstract

The Pairwise Sequentially Markov Coalescent (PSMC), and its extension PSMC′, model past population sizes from a single diploid genome. Both models have been widely applied, even to organisms with scaffold-level genome reference assemblies of limited contiguity. However it is unclear how PSMC and PSMC′ perform on short scaffolds. We evaluated `psmc` and `msmc`, implementations of the PSMC and PSMC′ models respectively, on simulated genomes with low contiguity, and compared results to those from fully contiguous data. Simulations with scaffolds from 100 Mb to 10 kb revealed that `psmc` maintains high accuracy down to lengths of 100 kb, while `msmc` is accurate down to 1 Mb. The discrepancy is not due to differing models, but stems from an implementation detail of `msmc`—homozygous tracts at the ends of scaffolds are discarded, making `msmc` unreliable for low contiguity genomes. We recommend excluding data that are aligned to shorter scaffolds when undertaking demographic inference.

## Introduction

The process of joining (coalescing) and splitting (recombining) lineages backwards-in-time for a sample of homologous sequences is described by the coalescent with recombination (Hudson, 1990). An important consequence of recombination is that there can be many distinct genealogies, known as marginal genealogies, at different locations along the sequence (Griffiths and Marjoram, 1997). The sequentially Markov coalescent (SMC, McVean and Cardin (2005)) models recombination as a Poisson process left-to-right along the sequence, approximating the coalescent with recombination by treating the marginal genealogy on the right of a recombination as a modification of the marginal genealogy on the left of the recombination. In this sense the approximation is a Markovian process along the sequence, and substantially reduces model complexity for long sequences compared to the full coalescent with recombination (Wiuf and Hein, 1999).

The Pairwise Sequentially Markov Coalescent (`PSMC`) uses a special case of the SMC approximation, restricted to pairs of sequences, to estimate the distribution of coalescent times within a single diploid genome (Li and Durbin, 2011). `PSMC` scans along a contiguous segment of the genome and considers marginal genealogies, using their distinct pairwise coalescent times as the unknown states in a hidden Markov model (HMM). To enable parameter estimation, continuous time is approximated by a finite partition of time intervals, and transition probabilities are inferred by Baum-Welch iteration of the forward-backward algorithm. Each genotype at consecutive genomic coordinates provides a new observation for the HMM, a homozygote or a heterozygote, with their emission probabilities determined by the pairwise coalescent time at the current locus, and the genome-wide mutation rate. The population size in a given time interval is inversely proportional to the rate of coalescence, as inferred by maximising the fit of the model to both the HMM transition matrix and the emission probabilities.

The Multiple Sequential Markov Coalescent (`MSMC`, Schiffels and Durbin (2014)) is an extension to `PSMC`, and models the distribution of first-coalescent times of two or more haploid sequences. If used with only two haploid sequences, `MSMC` closely matches the `PSMC` model, with the exception that it implements SMC′ (Marjoram and Wall, 2006), a refinement of SMC incorporating recombinations that immediately coalesce back to the same lineage. For this reason the `MSMC` model, when applied to a diploid genome, is referred to as `PSMC`′. Compared to `PSMC`, the genome wide recombination rate is more accurately estimated under the `PSMC`′ model, but population size estimates are qualitatively similar (Schiffels and Durbin, 2014).

Other approaches for inferring population size histories typically require either phased genotypes, multiple individuals, or both (Dutheil *et al.*, 2009; Gutenkunst *et al.*, 2009; Sheehan *et al.*, 2013; Boitard *et al.*, 2016; Terhorst *et al.*, 2017). However, in small scale studies of non-model organisms, it is common for only one individual, or a few individuals, from a single population to be sequenced, and genotypes are unlikely to be phased. Population size history, particularly in the recent past, can also be estimated from the length distribution of tracts of identity-by-descent (Palamara *et al.*, 2012), identity-by-state (Harris and Nielsen, 2013), or runs of homozygosity (MacLeod *et al.*, 2013). While potentially useful for a single diploid individual, such approaches are not readily applicable to short scaffolds, where such tracts may be broken across scaffold boundaries. In contrast, `PSMC` and `PSMC`′ are very attractive as they require only diploid genotypes for a single individual, which need not be phased.

By using the sequentially Markovian approximation, `PSMC` and derived methods implicitly assume that genomic information is contiguous. While initially applied to human datasets, which have very high contiguity, `PSMC` and `PSMC`′ have since been applied to many non-model organisms where the contiguity of genomic sequences may be poor (Zhao *et al.*, 2013; Dobrynin *et al.*, 2015; Mays *et al.*, 2018; Kozma *et al.*, 2016; Feigin *et al.*, 2018). In particular, demographic history is regularly inferred from a *de novo* assembly as part of genome sequencing projects. Due to time and funding constraints, genome assemblies are often constructed from only short read sequencing data, and assembled into contigs or short scaffolds. These cannot be ordered or oriented with respect to one another (violating the SMC model), nor anchored to physical chromosomes. Where sequencing data is aligned to such assemblies, the genomic information used for population size inference inherits the low contiguity of the assembly. While small gaps in coverage along a scaffold can be handled gracefully, the HMM must be applied separately to each distinct scaffold, and it is not clear what the length threshold is to obtain robust population size inferences.

## Results and Discussion

### Simulations

To assess the impact of reference genome contiguity on population size estimates, we simulated genomes for populations with three different demographic histories: a constant population size; a bottleneck; and recovery following a bottleneck (Fig. 1A). For each demographic scenario, we simulated 10 independent populations and sampled 20 *×* 100 Mb haploid chromosomes, representing 10 diploid genomes from each population. New datasets were then created by fragmenting each genome into equally sized scaffolds at four distinct lengths, 10 Mb, 1 Mb, 100 kb, and 10 kb. Population size histories were then inferred for all fragmented and unfragmented datasets using `psmc` (Li and Durbin, 2011) and `msmc` (Schiffels and Durbin, 2014), implementations of `PSMC` and `PSMC`′ respectively.

**Figure 1:**
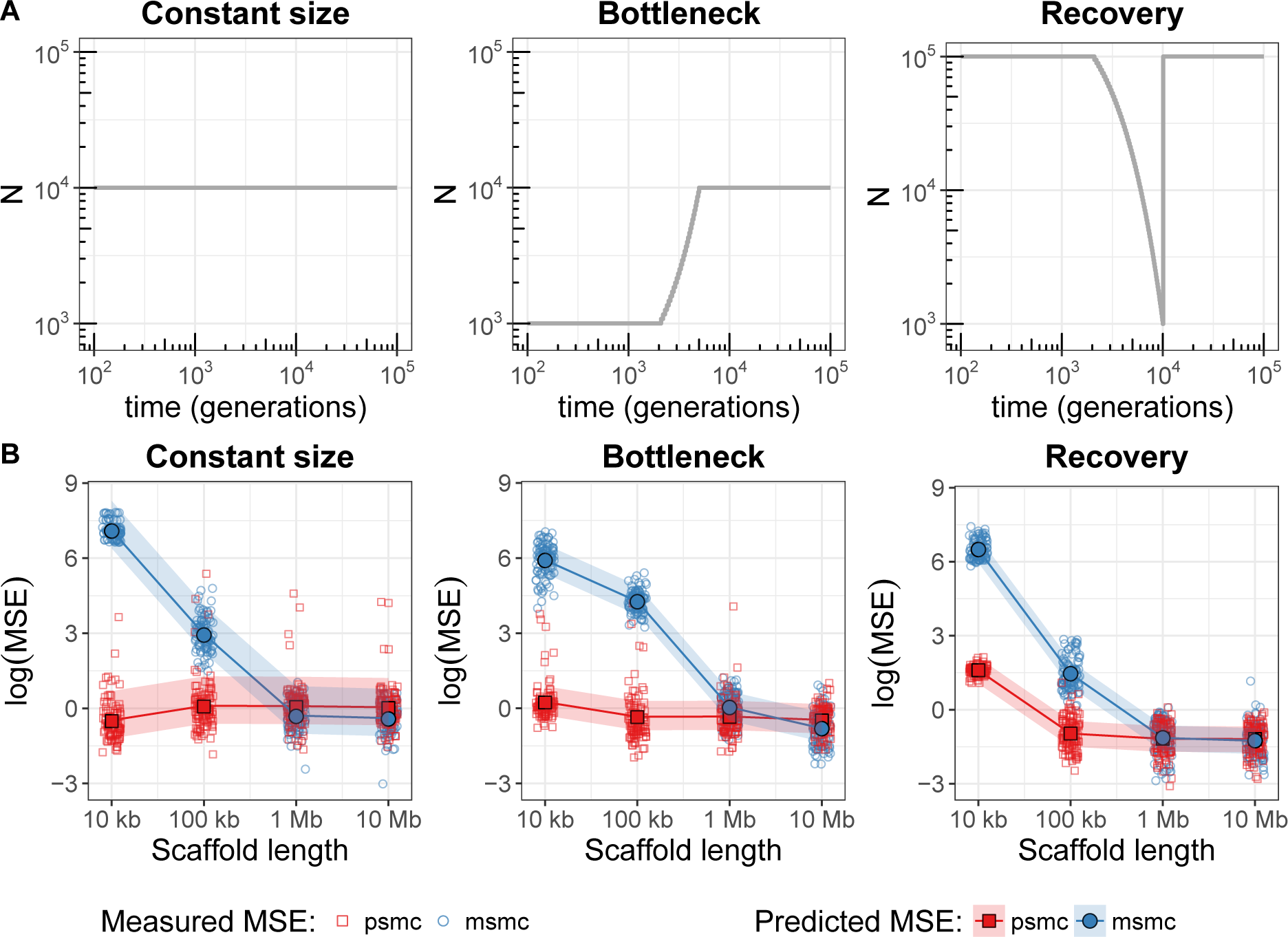
**A)** Simulated population size histories. **B)** Mean squared error (MSE) of population size inferences from simulations shown immediately above. Larger values indicate a loss of accuracy in the population size estimate. Small hollow markers indicate MSE for distinct simulated individuals (100 Mb per individual; 10 individuals each from 10 populations), with red squares for `psmc` and blue circles for `msmc`. Data from each simulated individual was artificially fragmented to emulate genome sequences aligned to a scaffold-level reference assembly. At each scaffold length, MSE was calculated by comparing to inferences from unfragmented (100 Mb) scaffolds (see methods). Large solid markers and lines show predicted MSE from a linear mixed effect model, with 95% prediction intervals based on simulation.

### Mean squared error

In measuring the error of estimates, Li and Durbin (2011) compared population size inferences to the values that were simulated, but excluded time intervals in the recent and distant past. Population size estimates are expected to be unreliable for times outside a certain range since a typical genome contains relatively few breakpoints corresponding to recombination events in the very recent or very distant past. However, excluding temporal intervals requires advance knowledge of where the method may lose resolution, and this is dependent upon the population size history itself.

To quantify estimation error, we used inferences from the unfragmented datasets as the ‘truth’, not the values that were simulated. A loess smooth function (Cleveland *et al.*, 1992) was fitted to the unfragmented inferences for each simulated population, separately for `psmc` and `msmc`, using population size estimates from all individuals in a given population. Then for each simulated individual, the mean squared error (MSE) was measured between estimates from the fragmented datasets and the loess function for the corresponding population. The MSE was weighted, in discrete time intervals, using the inverse of the sample variance in estimates from the unfragmented datasets (the same individuals as used for the loess fit). This was done to avoid measuring error caused by limited genomic information about the recent and ancient past.

Comparisons of the MSE at each fragmentation level (Fig. 1B) suggest that shorter scaffolds do indeed result in less accurate population size estimates. Qualitatively, `msmc` appears to decline in accuracy at scaffold lengths between 1 Mb and 100 kb for all demographic scenarios, whereas `psmc` declines in accuracy only in the Recovery scenario, at scaffold lengths between 100 kb and 10 kb.

### Mixed effects model

To determine if the observed differences were significant, we fitted a linear mixed-effects model separately for each demographic scenario. The fixed effects were scaffold length and estimation program (`psmc` vs. `msmc`), and a random intercept was necessary to account for the repeated measures of each individual at multiple scaffold lengths. Both scaffold length and estimation program were found to be significant predictors of MSE in all demographic scenarios. Two-way interactions between scaffold length terms and estimation program were also significant in all scenarios.

### Empirical data

Arguably, the simulated population history scenarios are unrealistic. Simulated data also provides the best possible case in terms of missing data in that there is none. To gauge the impact of using a scaffold-level assembly with real data, we artificially fragmented chromosome 1 from a high coverage human genome, HG00419, a Southern Han Chinese female (The 1000 Genomes Project Consortium, 2015). Population size histories were again estimated using `psmc` and `msmc`, for each of the fragmented and unfragmented datasets (Fig. 2).

**Figure 2:**
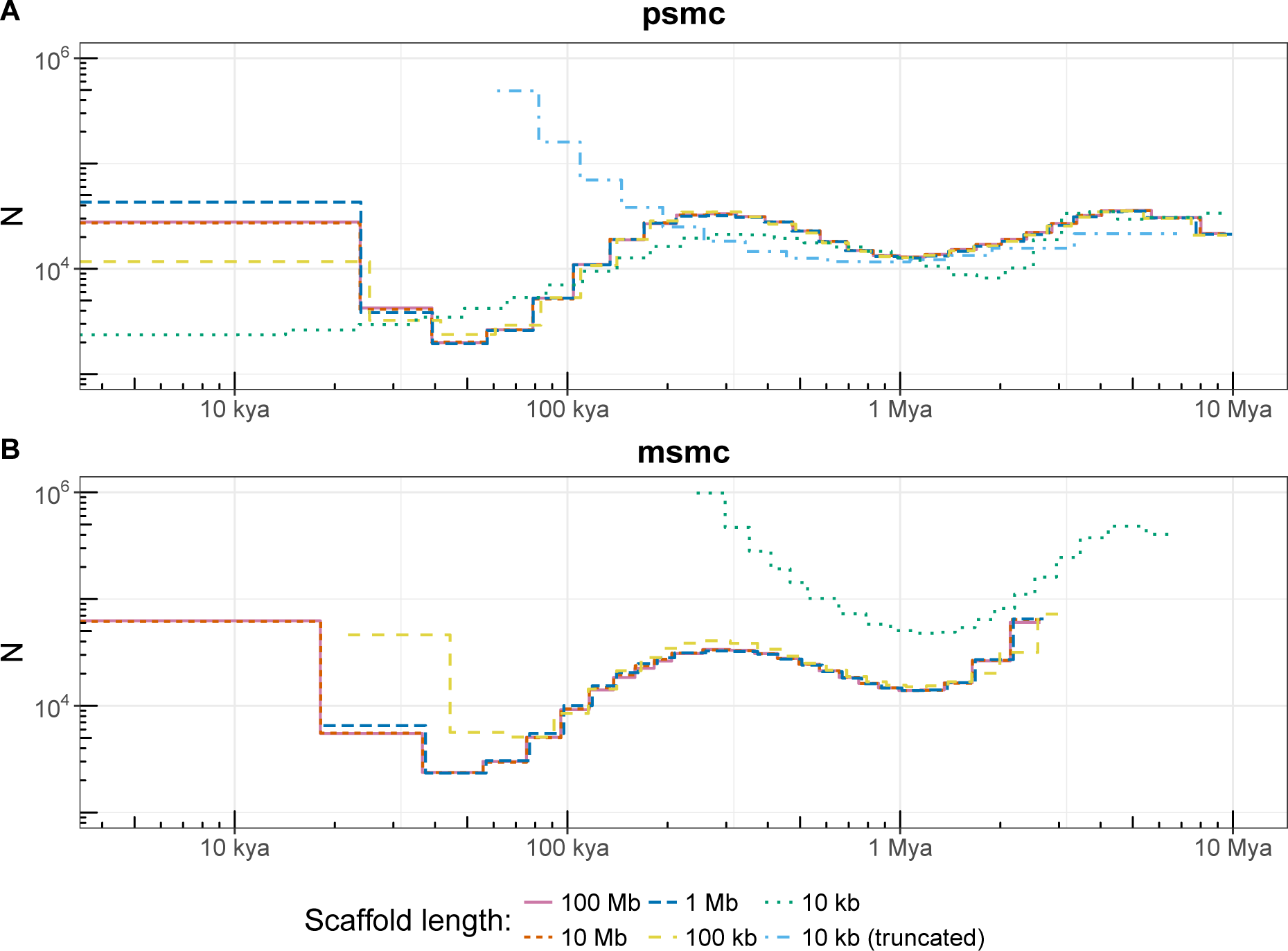
Population size history of HG00419, a Southern Han Chinese individual (The 1000 Genomes Project Consortium, 2015), inferred by **A)** `psmc` and **B)** `msmc`. Empirical data was artificially fragmented to emulate genome sequences aligned to scaffold-level reference assemblies. Population size inferences from `psmc` are consistent down to 100 kb scaffold lengths, with loss of resolution at 10 kb. For `msmc`, stable inferences can be made down to 1 Mb, but accuracy at 100 kb is poor in the recent past, and at 10 kb even broad demographic trends are difficult to discern. Input data to `psmc` for the ‘10 kb (truncated)’ line style had trailing homozygous sites removed from all scaffolds, to match the information content of `msmc` input. Plots were scaled to real time using a 25 year generation time and 1.25*e −* 8 mutations per base per generation. kya: thousand years ago; Mya: million years ago;

Both programs produced largely the same demographic history when processing long scaffolds, although `msmc` did not estimate population sizes for time intervals as far into the past as `psmc` (3 Mya vs. 10 Mya). For 10 kb scaffold lengths, inferences from `msmc` are substantially different to those using longer scaffolds, and a small departure is also discernible in the recent past for 100 kb scaffolds. Estimates from `psmc` have noticeably poorer resolution at the 10 kb scaffold length, but are remarkably consistent for longer scaffolds.

The data conversion script provided with `psmc` (fq2`psmc`fa) ignores scaffolds having fewer than 10000 genotype calls by default. This excluded most of the 10 kb scaffolds, due to the presence of one or more missing genotypes. Disabling this filter to retain all scaffolds only marginally improved population size estimates, and only in more ancient time intervals (results not shown). We considered the possibility that with 10 kb scaffolds, `psmc` might still accurately recapitulate the population size history if provided with more information. To this end chromosome 2 was also partitioned into 10 kb scaffolds and appended to the chromosome 1 data (doubling the information to *∼*500 Mb in total). However, the additional information did not alter the result.

### msmc discards homozygous tracts at the ends of scaffolds

An input file for `msmc` contains lines that specify the coordinate of a heterozygote site and its distance from the previous heterozygote on the same scaffold. Nothing is specified for coordinates after the last heterozygote, and the scaffold is implicitly truncated here. For short scaffolds this causes substantial information loss. Indeed, short scaffolds may contain no heterozygote sites at all, and input files for such scaffolds are empty.

To determine if truncation was a major cause of the different behaviour between `psmc` and `msmc`, we ran `psmc` on 10 kb scaffolds that were artificially truncated to match the information available to `msmc`. Scaffolds containing no heterozygotes were omitted. This output (‘10 kb (truncated)’ in Fig. 2A), shows a similar trend to that for `msmc` on 10 kb scaffolds, although differences remain.

Marginal genealogies with recent coalescent times have accumulated few mutations, so corresponding regions of the genome contain mostly homozygote genotypes. Truncation increases the proportion of heterozygotes, hence recent coalescent times appear older. On short scaffolds, all marginal genealogies are near a scaffold end, so inferences from short truncated scaffolds are more strongly biased to not observe recent coalescent events. Since the population size for each time interval is inversely related to the rate at which pairs of haplotypes coalesce, the smaller number of observations of high homozygosity genomic tracts also means that population size inferences are biased upwards. Both artefacts are noticeable, particularly in the more recent time bins, for `psmc` with artificially truncated 10 kb scaffolds (Fig. 2A) and for `msmc` with 10 kb and 100 kb scaffolds (Fig. 2B).

## Conclusion

Reasonable parameter inference in a hidden Markov model relies on observations leading up to, and following, transitions in state. For `PSMC`, this corresponds to having sufficient sequence contiguity to observe genomic tracts on both sides of historical recombination breakpoints. The chance that a short scaffold will contain a tract covering a recombination breakpoint depends not only on the completeness of the reference assembly, but also the sparsity of breakpoints.

Several factors contribute to breakpoint density, including population size, the per base recombination rate, and recombination hotspots. A population suffering a recent and very severe bottleneck will give rise to mostly recent pairwise coalescent times, and few recombination breakpoints, both of which are poorly represented within short scaffolds. Our simulations considered a mammalian recombination rate

(3.125 × 10^−9^ per base per generation) and population size histories that are relevant to many taxa. This >suggests that `PSMC` inference can be reasonable from scaffolds as short as 100 kb for a wide range of datasets.

Scaffold level reference assemblies are unlikely to contain equally sized scaffolds, as evaluated here. Generally, a scaffold-level assembly contains tens of long scaffolds and tens of thousands of short scaffolds.

In such cases, it is reasonable to exclude scaffolds shorter than 100 kb when running `psmc`, and scaffolds shorter than 1 Mb for use with `msmc`. However, we caution that this guideline may be too optimistic for severely bottlenecked populations or genomic data aligned to a very low quality reference assembly.

## Materials and Methods

### Simulations

Simulations were performed using scrm (Staab *et al.*, 2015), with mutation rate *µ* = 1.25 *×* 10^*−*8^ per base per generation and recombination rate *µ/*4 per base per generation. Simulation output was artificially fragmented during conversion to `psmc` and `msmc` input formats, using a custom Perl script. Demographic inferences were obtained from `psmc` v0.6.5-r67 and `msmc` v1.0.0 for all inputs. Both `psmc` and `msmc` were run with the same time bin parameter (-p 1*2+15*1+1*2), although we note that each program calculates time boundaries for the discrete bins differently, so a completely fair comparison is not possible. Scripts used for simulation, format conversion, and running `psmc`/`msmc` are available from https://github.com/grahamgower/psmc-error-analysis/.

### Mean squared error

For each simulated population history scenario and each estimation program, estimates from the unfragmented datasets were used to fit a loess function of log population (log(*N*)) against log time (log(*t* + 10)). The offset of 10 was based on a sensitivity analysis and the smallest non-zero time. An optimal value for the loess smoothing parameter was selected by maximising the corrected AIC (AICc) (Hurvich *et al.*, 1998). Mean squared error for individual *i* in population *j* was calculated as

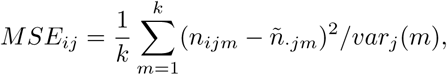

where the sum extends over all *k* time intervals, *n_ijm_* is the log of the population size estimate in interval *m*, and *ñ_⋅jm_* is the prediction for the *m*th time interval from the loess function fitted for the *j*th population. The variance step function *var_j_*(*m*) at time interval *m*, for the *j*th population, was calculated by splitting time on a log scale into 10 even-width bins and calculating the variance in each bin.

### Mixed effects modelling

Scatter-plots of MSE against scaffold length indicated a cubic relationship between MSE and log(scaffold length). This was confirmed by comparing residual plots for linear, quadratic, and cubic models. To help numerical consistency of the fitting process, we performed a location scaling of log(scaffold length).

Bivariate analysis of each of the predictors—log(scaffold length), estimation program, population history scenario, sample ID, and population ID—were used for variable selection. Only log(scaffold length), estimation program, and population history scenario had a significant relationship with MSE.

The linear mixed effects model was fitted using the lme4 package (Bates *et al.*, 2015) in R (R Core Team, 2017). The fixed effects were log(scaffold length) and estimation program. Up to two-way interaction terms were considered for each of the cubic log(scaffold length) terms with estimation program. To account for repeated measures from each simulated individual due to multiple levels of fragmentation, we included random effects. Both random intercepts and random slopes were considered.

All significance testing was performed using the lmerTest package (Kuznetsova *et al.*, 2017). All assumptions of the linear mixed-effects models were assessed and regarded as reasonable. The 95% prediction intervals were based on simulation with the merTools package (Knowles and Frederick, 2016).

### Empirical dataset

We downloaded the cram alignment file for HG00419, aligned to assembly GRCh38DH, from The 1000 Genomes ftp server, and called genotypes with samtools -q20 -Q20 -C50 … | bcftools call-c …. The resulting vcf was partitioned into scaffolds of a specific size by modifying the chromosome name and position to which each genotype call corresponded, and was performed separately for each of the scaffold sizes 100 Mb, 10 Mb, 1 Mb, 100 kb, and 10 kb. Input for both `psmc` and `msmc` were filtered to exclude sites with less than half, or greater than double, the mean depth (54.76). The vcf was converted to `psmc` input format with vcfutils.pl (distributed with samtools) and fq2`psmc`fa (distributed with `psmc`), then `psmc` was run with time bin parameter -p 4+25*2+4+6. The same vcf was converted to `msmc` input format with bamCaller.py and generate multihetsep.py, both distributed with `msmc`-tools, then `msmc` was run with parameters -R -p 15*1+15*2. The time bin parameters for both programs were chosen to be suitable for inferring human demography (Li and Durbin, 2011; Schiffels and Durbin, 2014).

## Acknowledgements

This work was supported by an Australian Government Research Training Program Scholarship [GG], and research fellowships from the University of Adelaide and the Australian Research Council [BL].

GG, JT, ABR, JS, BL, and NB designed the study. GG performed simulations and processed the empirical data. JT calculated the MSE and performed mixed effects modelling. All authors interpreted the results. GG wrote the manuscript with the help of all coauthors.

